# Multiplex Imaging Reveals Novel Subcellular, Microenvironmental, and Racial Patterns of MRTFA/B Activation in Invasive Breast Cancers and Metastases

**DOI:** 10.1101/2024.01.03.573909

**Authors:** Stephanie M. Wilk, Kihak Lee, Alexa M. Gajda, Mohamed Haloul, Virgilia Macias, Elizabeth L. Wiley, Zhengjia Chen, Xinyi Liu, Xiaowei Wang, Maria Sverdlov, Kent F. Hoskins, Ekrem Emrah Er

**Affiliations:** Department of Physiology and Biophysics, College of Medicine, University of Illinois Chicago, Chicago, IL; Department of Pathology, University of Illinois Chicago, Chicago, IL; Division of Epidemiology and Biostatistics, School of Public Health, University of Illinois Chicago, Chicago, IL; Biostatistics Shared Resource, University of Illinois Cancer Center, Chicago, IL; Department of Pharmacology & Regenerative Medicine, College of Medicine, University of Illinois Chicago, Chicago, IL; Research Histology Core, Research Resources Center, College of Medicine, University of Illinois Chicago, Chicago, IL; Division of Hematology/Oncology, College of Medicine, University of Illinois Chicago, Chicago, IL

**Keywords:** Myocardin related transcription factors, MRTFA, MRTFB, breast cancer, metastasis, tumor microenvironment, cancer health disparities, antigen presenting cells, dendritic cells, tumor immunity, immune checkpoint, VSIR

## Abstract

Breast cancer progression and metastasis involve the action of multiple transcription factors in tumors and in the cells of the tumor microenvironment (TME) and understanding how these transcription factors are coordinated can guide novel therapeutic strategies. Myocardin related transcription factors A and B (MRTFA/B) are two related transcription factors that redundantly control cancer cell invasion and metastasis in mouse models of breast cancer, but their roles in human cancer are incompletely understood. Here, we used a combination of multiplexed immunofluorescence and bioinformatics analyses to show that MRTFA/B are concurrently activated in tumor cells, but they show distinct patterns of expression across different histological subtypes and in the TME. Importantly, MRTFA expression was elevated in metastatic tumors of African American patients, who disproportionately die from breast cancer. Interestingly, in contrast to publicly available mRNA expression data, MRTFA was similarly expressed across estrogen receptor (ER) positive and negative breast tumors, while MRTFB expression was highest in ER+ breast tumors. Furthermore, MRTFA was specifically expressed in the perivascular antigen presenting cells (APCs) and its expression correlated with the expression of the immune checkpoint protein V-set immunoregulatory receptor (VSIR). These results provide unique insights into how MRTFA and MRTFB can promote metastasis in human cancer, into the racial disparities of their expression patterns, and their function within the complex breast cancer TME.

**Graphical Abstract:** 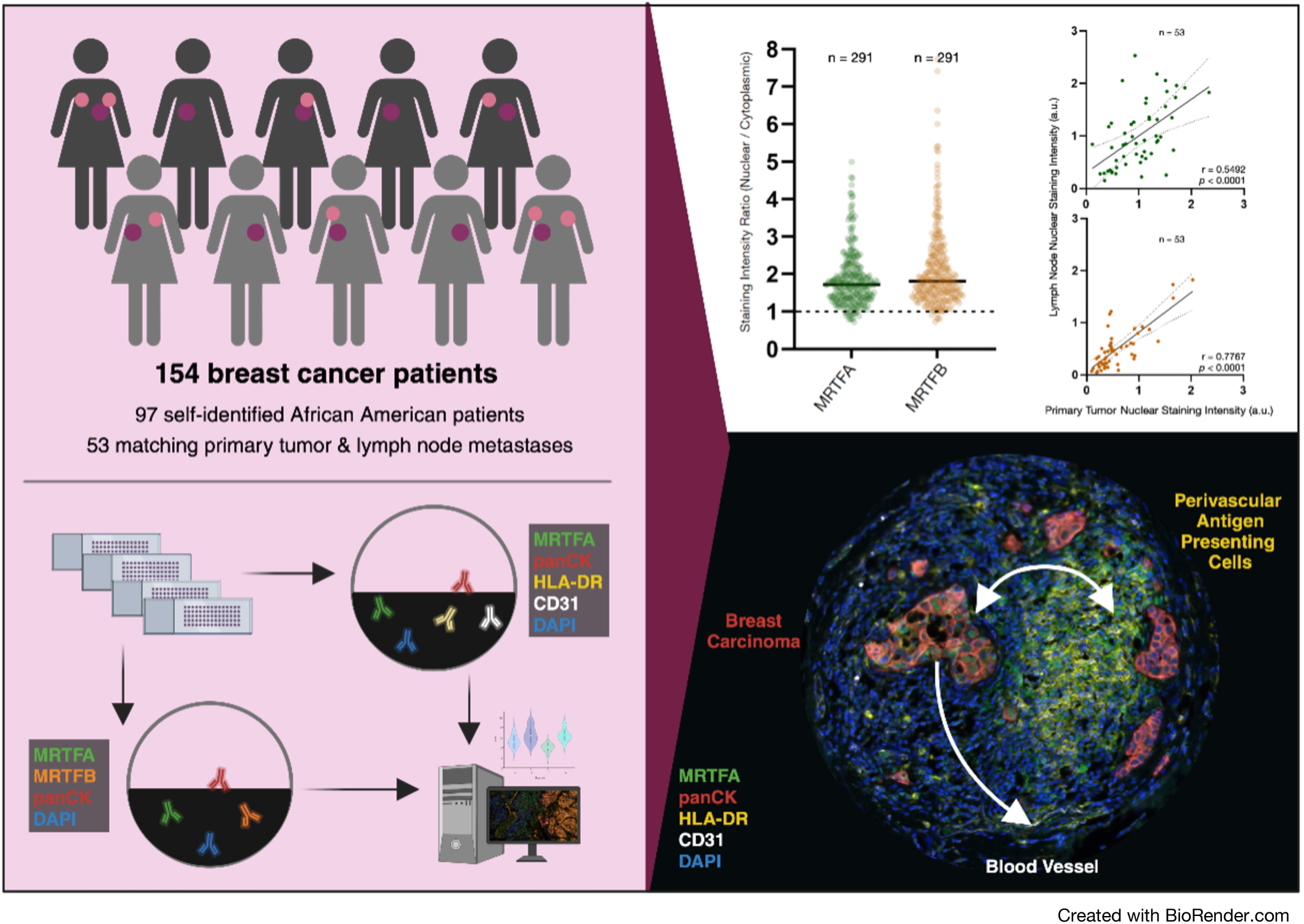

## Introduction

Breast cancer claims the lives of over 43,000 women in the United States each year and the vast majority of breast cancer-related deaths result from distant metastases^1,2^. Unfortunately, there are very few treatment options for women diagnosed with metastatic breast cancer and some existing therapeutic strategies are even less effective in prolonging the lives of vulnerable patient populations, such as African American women, who are more likely to die from breast cancer than women from other racial groups ^1,3,4^. This survival disparity is caused by structural racism and social determinants of health, which are thought to foster a chronic level of elevated stress that, in turn, influences tumor evolution, promotes an inflammatory microenvironment and leads to metastases^5–7^. Therefore, to create more equitable treatment strategies, it is imperative to identify molecular mechanisms of breast tumor initiation, progression, and metastasis in women from diverse racial and ethnic backgrounds.

Breast cancer is thought to evolve through multiple steps that start with ductal carcinoma *in situ* (DCIS), followed by progression into invasive breast cancer (IBC) and ultimately into metastases to lymph nodes and to vital secondary organs such as the bones, lungs, liver and the brain^8^. DCIS represents 20% of new breast cancer diagnoses and, unlike IBC, it rarely relapses as distant metastases following local treatment. DCIS involves a pre-invasive lesion of tumor cells isolated from the stroma via a barrier of myoepithelium and basement membrane proteins. In IBC, the myoepithelial layer is loosened and lost and the tumor cells overcome the stromal barriers to invade into the lymphovascular system for metastasis^9^. Recent studies seeking to understand the changes in the tumor microenvironment (TME) that accompany the transition from DCIS to IBC suggest that the structure of the tumor stroma can predict IBC relapse^10^. For example, DCIS cases that present with a thick, continuous E-cadherin-rich myoepithelium and accumulation of perivascular antigen presenting cells (APCs) are more likely to progress into IBC and relapse^10^. These collective changes are coordinated through precise action of transcription factors (TFs) that promote a tumor conducive TME and that regulate breast cancer cells’ proliferation, migration, invasion, survival at distant secondary organs, and evasion of the immune system^11^.

Myocardin related transcription factors A and B (MRTFA/B) are two related TFs that have been extensively studied for their pro-metastatic roles in experimental mouse models of breast cancer. MRTFA/B bind to the serum response factor (SRF), which recognizes CArG box DNA sequences, and their loss-of-function reduces actin cytoskeleton polymerization, perturbs cellular migration and contractility, and thereby inhibits subsequent metastatic invasion^12–15^. However, recent work showed that these pro-metastatic functions of MRTFA/B highly depend on the composition of the TME^16^. For example, despite their higher metastatic potential in immune compromised mice, MRTFA/B expressing cancer cells are eliminated by mechanosurveillance if the host has intact cytotoxic lymphocyte activity. Mouse genetics also highlights the potential roles of MRTFA/B in regulating cells of the TME, such as myoepithelial cells, endothelial cells, immune cells, and fibroblasts^9^. For example, MRTFA knockout results in lactation/involution defects in postpartum mice, suggesting that MRTFA is critical for myoepithelial function^17^. MRTFB knockout in mice is embryonic lethal because of defects in smooth muscle investment during cardiovascular development and MRTFA, MRTFB and SRF have all been shown to be critical for endothelial cell function during hemostasis and vascular remodeling^18–21^. In mouse models of breast tumorigenesis, MRTFA/B have been shown to enforce the pathological role of cancer associated fibroblasts (CAFs), which promote extracellular matrix deposition and remodeling to facilitate cancer cell invasion^22^. These findings highlight the importance of gaining a holistic view of the tumors and the TME when studying the functional relevance of tumorigenic and metastatic TFs, but despite these fundamental findings from mouse genetics, the tumor intrinsic and microenvironmental roles of MRTFA and MRTFB in human breast cancer remain incompletely understood. To address this, we decided to investigate the clinical, demographic, tumor-intrinsic and microenvironmental patterns of MRTFA/B’s expression and activation by using a combination of multiplex imaging on multiple racially diverse tissue microarrays (TMA) and bioinformatics analyses of The Cancer Genome Atlas (TCGA) and Molecular Taxonomy of Breast Cancer International Consortium (METABRIC) datasets.

## Materials and Methods

### Patient cohort

Demographic and histological summary of the UIC patient cohort is provided in **Table 1**, and this TMA includes breast cancer tissues from 97 Black/African American, 46 White, and 11 Other/Race Unknown self-identified women with a median age at diagnosis of 55 years (range 28-89 years). Invasive ductal carcinoma is the predominant histological subtype (n = 118). Half of cases involve high-grade tumors (n = 77). Clinical immunohistochemistry (IHC) and fluorescent in situ hybridization (FISH) results were used to determine estrogen receptor (ER), progesterone receptor (PR), HER2, Ki67, and p53 status of tumor samples. 68.1% of women present with ER+ tumors (n = 105). All samples were collected according to approved Institutional Review Board (IRB, UIC IRB Protocol # 2017-0466) protocols and patient information was de-identified.

**Table 1.**
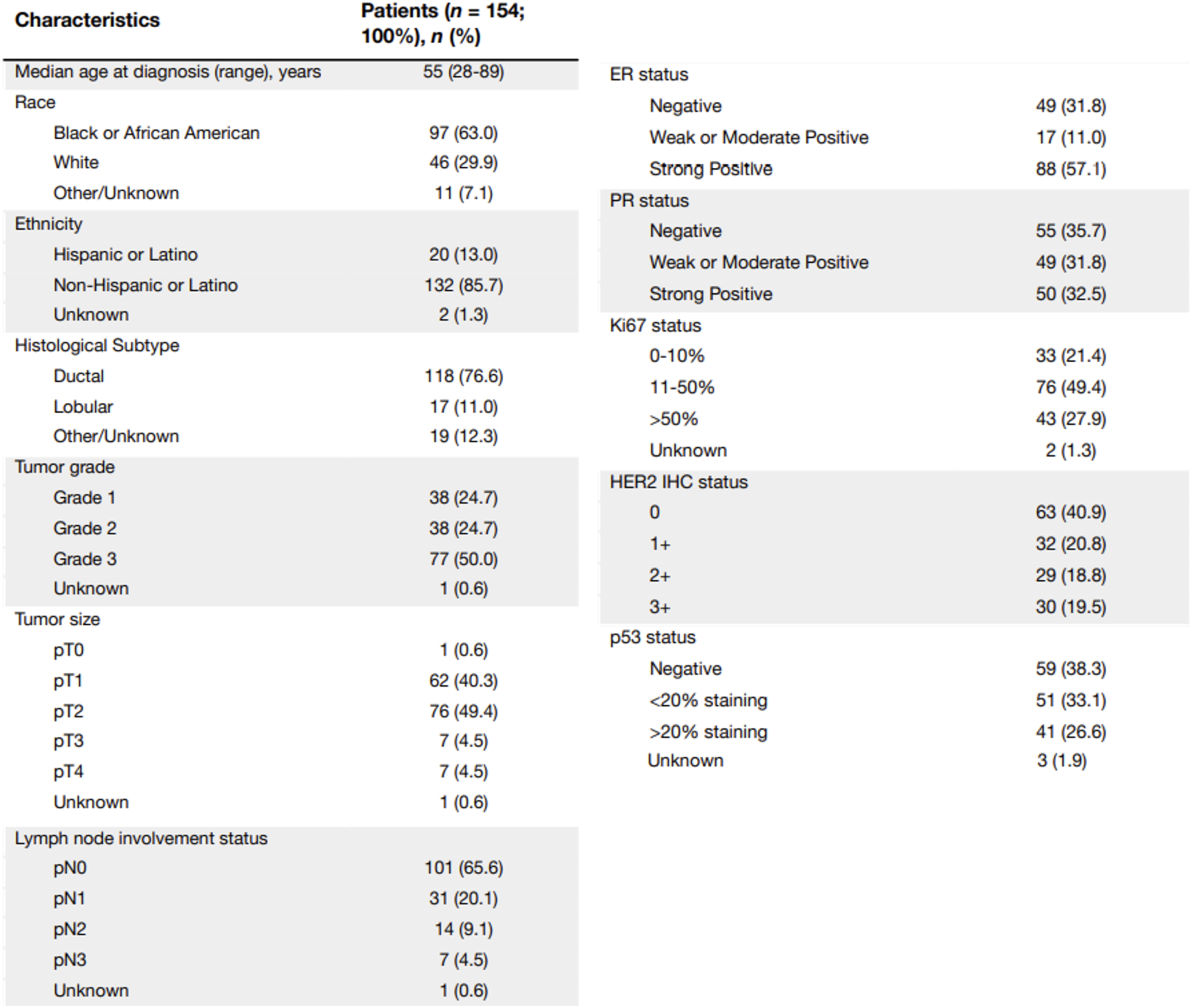
Patient characteristics.

### Breast tumor TMA construction

The UIC TMA (BRWG UIC-001-TMA) used in this study includes a consecutive series of invasive breast cancer cases undergoing surgery at the University of Illinois Cancer Center between 2013 and 2019 and is offered through the University of Illinois Cancer Center Breast Cancer Working Group (BCWG). Cases were arranged on 4 blocks, with many cases having matching normal breast tissue and/or lymph node metastases samples. Tumor cores (1.0 mm diameter) were sampled from areas of largest grade and focus by a breast pathology specialist. Most tumor samples are arrayed in duplicate, and some have cores from additional tumor foci. All normal breast tissue and lymph node metastases cores are arrayed in singlicate. Cases with neoadjuvant therapy were excluded from this TMA.

### Multiplex immunofluorescence staining

We optimized a six-color Opal TSA multiplex immunofluorescence panel labeling MRTFA, MRFTB, CD31, HLA-DRA, panCK, and the nuclear marker DAPI on breast cancer control samples. Key elements of the protocol are described in **Table 2**. Staining was performed on a Leica BOND RX autostainer according to the modified Opal 7-color (v5.2 plus) preset protocol using the Opal 6-Plex Detection kit for Whole Slide Imaging (Akoya Biosciences). Prior to the first staining round, tissue was subjected to sequential heat-based antigen retrieval for 40 minutes at 99°C with Bond Epitope retrieval buffer 2 (pH 9.0) and buffer 1 (pH 6.0). For each staining round, tissue sections were incubated with 1X Antibody Diluent/Block (Akoya Biosciences) for 5 minutes, followed by the incubation with primary antibody for 30 minutes, Opal Polymer-HRP for 10 minutes, and corresponding Opal dye for 10 minutes. All incubations were conducted at room temperature. After each round of staining, slides were incubated with Bond Epitope retrieval buffer 1 for 30 minutes at 98°C to remove primary and secondary antibodies. For optimization, serial dilutions of each antibody were run as single stains with the corresponding Opal dye detection, and signal localization and morphology were compared to those detected in single chromogen IHC staining performed on adjacent slide. The built-in spectral library was used for signal unmixing. All optimization images were unmixed and assessed in inForm V2.6 and Phenochart V1.1.0 (Akoya Biosciences).

**Table 2.** Multiplex immunofluorescence staining protocol.

|  | Antibody | Vendor | Catalog # | Clone | Conc. | Opal dye | Opal dilution |
| --- | --- | --- | --- | --- | --- | --- | --- |
| 1 | MRFTA | Sigma-Aldrich | HPA030782 | polyclonal | 1:50 | 520 | 1:100 |
| 2 | MRTFB | Bethyl Laboratories | A302-768A | polyclonal | 1:50 | 620 | 1:150 |
| 3 | PanCK | Agilent DAKO | M3515 | AE1/AE3 | 1:200 | 690 | 1:300 |
| 4 | HLA-DR | Abcam | ab92511 | EPR3692 | 1:5000 | 570 | 1:150 |
| 5 | CD31 | Cell Signaling | 3528 | 89C2 | 1:2000 | 780 | 1:100/1:25 |

### Image analysis

Slides were scanned on PhenoImager HT (Akoya Biosciences) in 4-color motif mode. InForm software (Akoya Biosciences) was used to remove autofluorescence and cleanly separate signals from each target into individual channels. Image analysis was performed using HALO software (Indica Labs). Images were manually annotated to identify tissue and remove artifacts. HALO mininet AI classifier was trained to identify panCK-positive regions, representing tumor areas in tumor cores and epithelial areas in normal cores. This classifier was used to label each cell as either within or outside of the tumor/epithelial area. Nuclei were segmented via a custom-trained HALO AI nuclear segmentation classifier. Cytoplasm regions were approximated by segmenting a ring around each nucleus. Mean pixel intensity was reported for MRTFA and MRTFB in the nucleus, cytoplasm, and whole cell areas.

### Statistical analysis

Statistical analyses were performed using GraphPad Prism 9 (GraphPad Software Inc.). Analyses considered staining intensity values from the nuclear compartment, cytoplasmic compartment, and whole cell area of primary tumor and lymph node metastases cores. Cores that lacked staining intensity values for either compartment or the whole cell area due to factors such as loss of tissue or poor staining quality were excluded from analyses. Fewer than 20 tumors had a secondary tumor and thus secondary tumor cores, when available, were excluded. In the case of multiple primary tumor or lymph node metastasis samples from a single patient, the mean staining intensity values were calculated and used in the analyses. Primary tumor samples were matched to corresponding lymph node metastasis in 53 cases. Pearson’s correlation analysis and Wald’s test were performed to obtain and test the correlation coefficient values between the nuclear, cytoplasmic, and total cell expression of MRTFA/B in the individual tumor cells of primary tumor cores (**Fig. 2** and **Supplemental Fig. S1**). Pearson’s correlation analysis and Wald’s test for p-value, along with simple linear regression and a 99% confidence interval, were used to plot the staining intensities of matching primary tumor and lymph node metastasis cores (**Fig. 3A, 3C** and **Supplemental Fig. S2A-D).** Fisher’s exact test was used to explore how categorical clinical and demographic variables influenced the lymph node staining intensities of MRTFA/B (**Fig. 3B**), and continuous clinical and demographic variables were explored with multiple linear regression models (**Fig. 3D-G** and **Supplemental Fig. S2E-H**). Cases that did not have data available for every variable tested for were excluded from the analysis. Staining intensity values from our TMA and mRNA expression values from databases across hormone receptor types were plotted with Tukey-presentation box plots (**Fig. 4A-D, 4F**, **4G** and **Supplemental Fig. S3A, S3B**). CTL dysfunction, exclusion, and infiltration and MRTFA mRNA expression among African American and White American patients are shown with violin plots (**Fig. 6A-D**) and MRTFA staining intensity in stromal cells across racial groups and tissue types is shown with histograms (**Fig. 6E**). Histograms are plotted with mean and SEM error bars. Across all figures, the means between two groups means were compared with unpaired t-tests; in the case of non-Gaussian distribution, the Mann-Whitney test was used instead. For more than two groups, their means were compared with one-way ANOVA; in the case of non-Gaussian distribution, the Kruskal-Wallis test was used instead. A *P* value <0.05 was considered statistically significant.

### Bioinformatics analysis

Bulk RNA sequencing data for TCGA and METABRIC databases were accessed using the cBioportal. mRNA expression values for Tumor Immune Dysfunction and Exclusion analyses were downloaded from the Broad Institute and analyzed as previously described^38^. Single cell RNA sequencing data was accessed by the Broad Institute’s Single Cell Portal https://singlecell.broadinstitute.org/single_cell. Dendritic cell and cancer associated fibroblast inflitration scores were calculated by using the Tumor Immune Estimation Resource 2.0 web interface http://timer.cistrome.org/.

## Results

To measure the degree of MRTFA and MRTFB protein expression in human tumors, we worked with two breast cancer TMAs. The first TMA was from the Cooperative Human Tissue Network and included normal breast tissue, ductal carcinoma *in situ* (DCIS), invasive cancer and lymph node metastases samples (designated as CHTN_BrCaProg3). The second TMA was designed by the Breast Cancer Working Group at the University of Illinois Cancer Center to represent our Cancer Center’s patient population residing in neighborhoods on the West Side and South Side of Chicago (designated as BRWG UIC-001-TMA). In contrast to commercially available alternatives, this unique TMA has a strong representation of African American breast cancer patients: 97 out of 154 patients who donated their samples self-identified as African American (**Fig. 1A**). Almost all tumors had a matching uninvolved adjacent breast tissue sample and 53 patients had matching lymph node metastases, which allowed us to measure MRTFA and MRTFB expression across tumorigenesis and metastasis. We conducted the first round of multiplex imaging on the CHTN_BrCaProg3 TMA by using an MRTFA antibody, an MRTFB antibody, panCytokeratin epithelial marker (panCK) and, DAPI for nuclear identification and by using the Opal-TSA detection system (**Fig. 1B**). In the uninvolved breast tissue, MRTFA antibody strongly labeled the nucleus of panCK+ myoepithelial cells as judged by basal position of these cells with respect to the mammary alveoli, but MRTFB expression was mostly non-detectable (**Fig. 1C**). The strong MRTFA protein expression is consistent with mouse genetics studies, where MRTFA knockout most prominently impacts lactation/involution cycle^17^. Interestingly, we observed that in IBC samples, this strong MRTFA signal in the myoepithelial compartment was lost, which is consistent with the loss of myoepithelial layer during transition from DCIS to IBC (**Fig. 1D** and Ref 23). Interestingly, in multiple cores in the CHTN_BrCaProg3 TMA, we found that cancer cells gained MRTFA and MRTFB signal in primary tumor cells and lymph node metastases, yet there were also instances of DCIS and IBC where no MRTFA/B expression was detected. These data suggest that transformation and tumor progression is coupled to expression of MRTFA/B in a subset of breast tumors.

**Figure 1.**
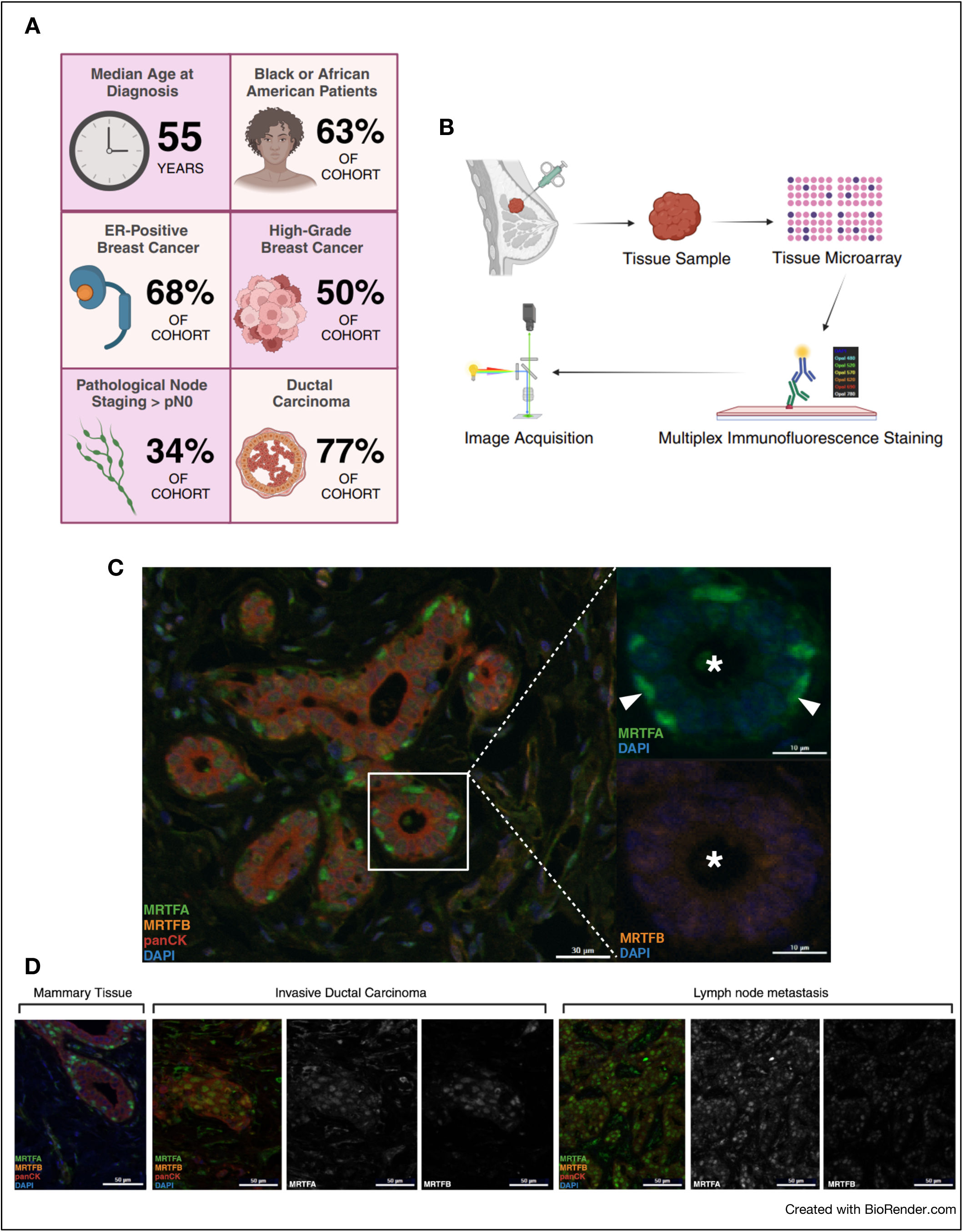
MRTFA and MRTFB show distinct expression patterns in the normal mammary gland. (A) Schematic representation of selected characteristics of patient cohort. (B) Illustrative summary of methods to image MRTFA and MRTFB in the tumor microenvironment. (C) Representative images of expression patterns of MRTFA and MRTFB in the normal mammary gland and its compartments: MRTFA (Opal 520) in green, MRTFB (Opal 620) in cyan, panCK (Opal 690) in red, and DAPI in blue. Asterisks (**\***) represent lumen of mammary epithelium and arrowheads (➤) point to the location of contractile myoepithelial cells. (D) Representative images of expression patterns of MRTFA and MRTFB in mammary tissue, Invasive Ductal Carcinoma, and lymph node metastases: MRTFA (Opal 520) in green, MRTFB (Opal 620) in cyan, panCK (Opal 690) in red, and DAPI in blue.

Next, we decided to measure the degree of MRTFA/B activation, which requires their nuclear localization. Both in 2-dimensional (2D) tissue culture experiments and in 3D models of mammary sphere formation, MRTFA/B nuclear localization is driven by acute stimuli, such as mitogens or cell adhesion to extracellular matrix, but nuclear MRTFA is exported to the cytoplasm by G-actin binding in asynchronously growing cells^15,24–26^. Therefore, we set out to measure MRTFA/B nuclear/cytoplasmic ratios as proxies for their activation state. We used the HALO AI neural network driven cell classification module for identifying panCK^+^ tumor cells and the nuclei segmentation module for cytoplasmic and nuclear signal detection in each cell in each of the primary tumors and metastatic lesions in the BRWG UIC-001-TMA (**Fig. 2A**). We found that in the vast majority of tumor cores the average MRTFA/B nuclear/cytoplasmic ratios were above 1, which suggests that MRTFA/B were more likely to be active when expressed (**Fig. 2B**). MRTFA and MRTFB perform redundant functions in tumorigenesis and metastasis in experimental models and we reasoned that if they performed redundant functions in human tumors, the expression of one MRTF would alleviate the need for the expression of the other^13,14,16^. Therefore, we calculated the Pearson correlation coefficient between MRTFA and MRTFB nuclear intensities in each cancer cell in each primary tumor core. We found that in 181 cores, Pearson correlation was higher than 0.5, while only 2 tumor cores had a negative correlation coefficient (**Fig. 2C, Supplemental Fig. S1A, S1B**) The average MRTFA nuclear signal intensity also positively correlated with the average MRTFB intensity (**Fig. 2D, Supplemental Fig. S1C, S1D**). This pattern of concurrent MRTFA and MRTFB expression suggests that MRTFA expression does not alleviate the need for MRTFB expression and that MRTFA and MRTFB can perform non-redundant functions in tumorigenesis and metastasis.

**Figure 2.**
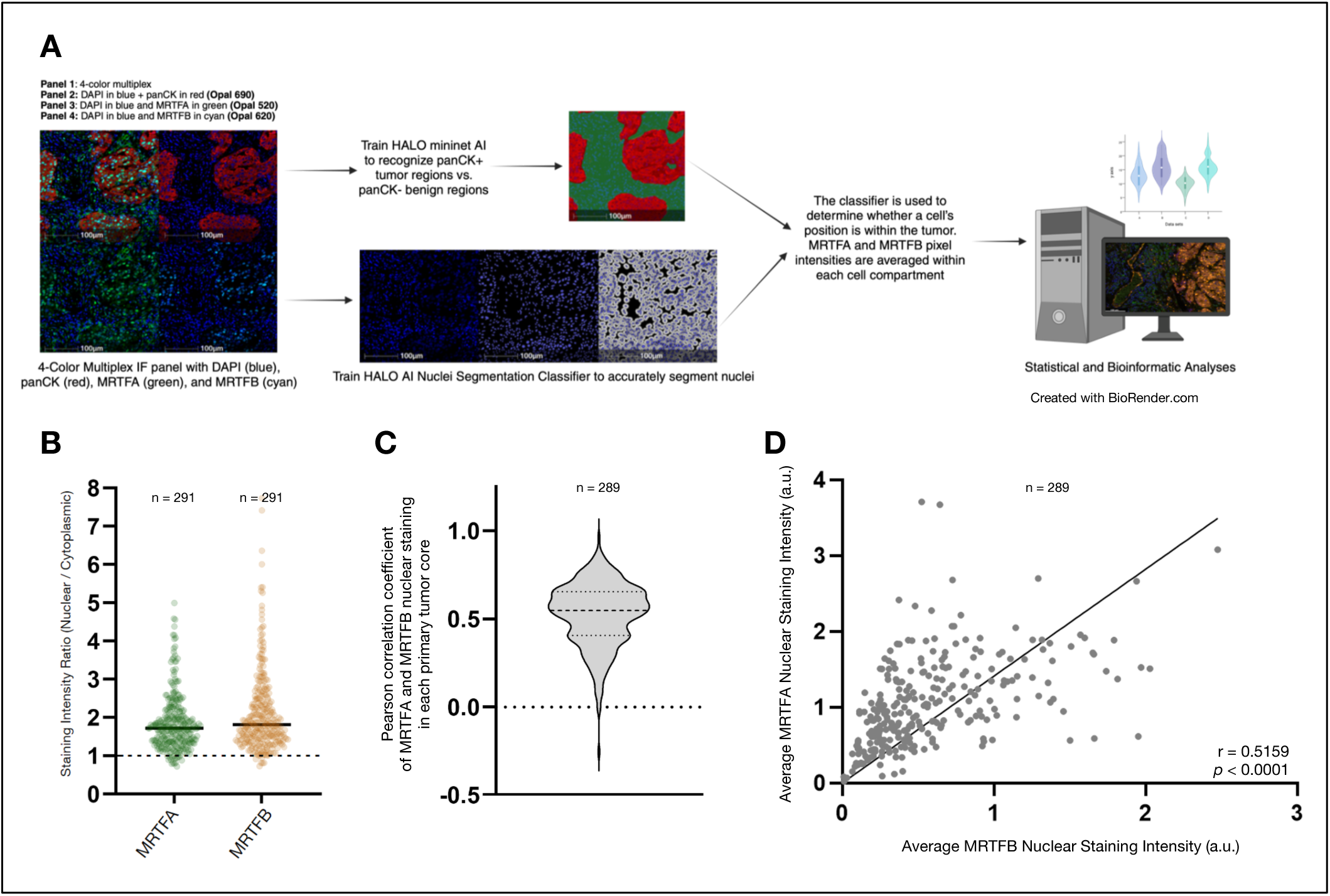
Subcellular segmentation reveals a correlation between MRTFA and MRTFB staining in tumor cells and abundant nuclear localization. (A) Workflow for nuclear and cytoplasmic segmentation of 4-color multiplex immunofluorescence in preparation for statistical and bioinformatic analyses, including training of HALO AI to recognize panCK-positive tumor regions versus panCK-negative benign regions of cores. (B) Violin plot of calculated staining intensity ratios (quotient of nuclear staining intensity and cytoplasmic staining intensity) for MRTFA and MRTFB in panCK-positive tumor cells of primary tumor cores (n = 291). Median staining intensity ratios greater than 1 suggest activation and abundant nuclear expression. (C) Violin plot showing distribution of Pearson correlation coefficients of average nuclear MRTFA and MRTFB staining intensities in panCK-positive tumor cells of primary tumor cores (n = 289). Mann-Whitney test used to calculate *p*-values. (D) Pearson correlation plot and simple linear regression of average nuclear MRTFA and MRTFB staining intensities in panCK-positive tumor cells of primary tumor cores (n = 289).

Metastatic cancer cells often retain and enrich for the expression of proteins that promote cancer cell invasion and dissemination as these proteins can provide a survival advantage during outgrowth at distant organs^11,27,28^. However, continued MRTFA/B activity can also pose an immune vulnerability at metastatic sites^14,16^. Therefore, to determine whether MRTFA/B expression was retained or lost at the secondary sites, we measured the average MRTFA/B staining intensities in lymph node metastases and compared it to their expression in primary tumors (**Fig. 3A**, and **Supplemental Fig S2A, S2B**). We found that lymph node staining intensity positively correlated with tumor staining intensity for both MRTFA and MRTFB. To explore how clinical and demographic parameters influenced these staining patterns, we ran a series of Fisher’s tests in which we dichotomized cases based on their MRTFA/B staining level with respect to the group’s median staining (**Fig. 3B**). Interestingly, of the 9 clinical and demographic parameters tested, self-identifying as African American was the only significant parameter for MRTFA lymph node staining intensity. In addition to race, having a progesterone receptor-positive (PR+) tumor was a significant parameter for MRTFB lymph node staining intensity. Since race was a shared significant parameter, we wanted to know if separating African American patients’ samples from White American patients’ samples would reveal distinct distribution patterns for matched primary tumor and lymph node metastasis cores (**Fig. 3C** and **Supplemental Fig. S2C, S2D**). Indeed, African American patients’ samples had noticeably steeper regression lines for both MRTFA and MRTFB and higher MRTFA lymph node intensities. Next, we ran multiple linear regression models for predicting MRTFA and MRTFB lymph node staining intensity values based on parametric and continuous variables such as age and primary staining intensity (**Fig. 3D-G** and **Supplemental Fig. S2E-S2H**). Consistent with the results of the Fisher’s tests, African American race significantly and positively correlated with MRTFA lymph node staining intensity (**Fig. 3E**). In contrast, the positive correlation between race and MRTFB staining intensity and PR status was lost (**Fig. 3G**). These results demonstrate that MRTFA and MRTFB expression levels are retained during the metastatic cascade and that the most significant parameter for MRTFA/B expression was race and likely the associated social determinants of health, but not the hormone receptor status.

**Figure 3.**
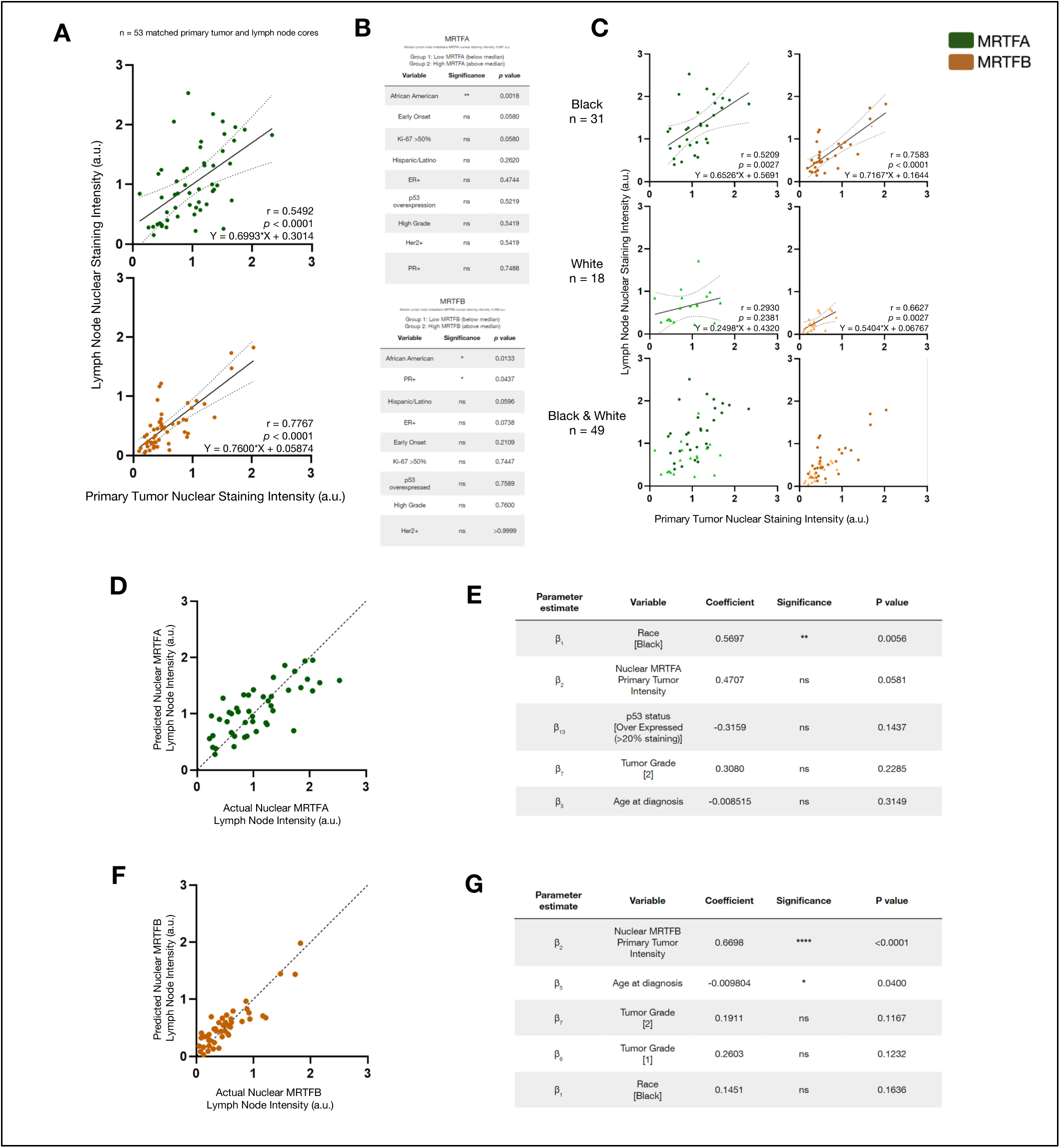
Clinical and demographic correlates of MRTFA and MRTFB expression in lymph node metastases. (A) Pearson correlation plots of primary tumor nuclear staining intensity and matching lymph node metastasis nuclear staining intensity of MRTFA and MRTFB for 53 cases. Dotted lines signify 99% confidence interval. Cases were categorized into “Low MRTFA/B” and “High MRTFA/B” groups based on falling above or below the median nuclear lymph node staining intensity. (B) Chart with results from a series of Fisher’s exact tests run for patient demographic and tumor characteristics variables. Cases that did not have data available for every variable tested for were excluded from the analysis (n = 8). (C) Pearson correlation plots of primary tumor nuclear staining intensity and matching lymph node nuclear staining intensity of MRTFA and MRTFB for self-identified Black/African American patients (n = 31) and White patients (n = 18). Dotted lines signify 99% confidence interval. (D) Multiple linear regression model for nuclear MRTFA lymph node intensity with patient demographic and tumor characteristics predictor variables. (E) Multiple linear regression results with top five predictor variables for nuclear MRTFA lymph node intensity based on *p*-value. (F) Multiple linear regression model for nuclear MRTFB lymph node intensity with patient demographic and tumor characteristics predictor variables. (G) Multiple linear regression results with top five predictor variables for nuclear MRTFB lymph node intensity based on *p*-value. Significance level for *p*-values: * (*p*<0.05), ** (*p*<0.01), *** (*p*<0.001), **** (*p*<0.0001).

The lack of evidence for a role of hormone receptor status in dictating MRTFA/B expression patterns was surprising because MRTFA expression is primarily thought to be prevalent in triple negative breast cancers and ER positive breast cancer models do not express high levels of endogenous MRTFA^29,30^. Therefore, we decided to compare the MRTFA/B expression levels across hormone receptor subtypes in TCGA and METABRIC cohorts to MRTFA/B expression levels in our BRWG-UIC-001 TMA. *MRTFB* mRNA expression and protein expression was significantly lower in basal and ER-negative tumors across all platforms (**Fig. 4A-4D and Supplemental Fig. S3A, S3B**). However, MRTFA mRNA and protein expression across different subtypes in our TMA, TCGA and the METABRIC samples were discordant: *MRTFA* mRNA was more highly abundantly expressed by the ER negative and basal cancer patient cohorts in the TCGA and METABRIC datasets, respectively, but MRTFA protein expression was lower in the BRWG-UIC-001 TMA ER negative primary tumor samples. We reasoned that the reason *MRTFA mRNA* is higher in the ER-negative samples in TCGA and METABRIC may be that ER-samples have a higher degree of stromal and immune cell content than ER+ samples and that these TME cells also express MRTFA^31^. To test this, we used the Tumor Immune Estimation Reporter (TIMER2.0) algorithm and calculated the tumor purity as a function of *MRTFA* (also known as *MKL1*) and *MRTFB* (*MKL2*) expression. Indeed, we found that MRTFA, but not MRTFB expression was negatively correlated with tumor purity (**Fig. 4E**). Furthermore, we found that in the BRWG-UIC-001 TMA, even though the MRTFA/B signals were significantly lower in stromal cells than in cancer cells, the stroma/cancer MRTFA signal ratio (∼0.81) was much higher than that of stroma/cancer MRTFB signal ratio (∼0.44) (**Fig. 4F, 4G**). Taken together, these results highlight the subtype specific expression patterns of MRTFA and MRTFB and reveal the TME as a significant source of MRTFA expression in breast cancers.

**Figure 4.**
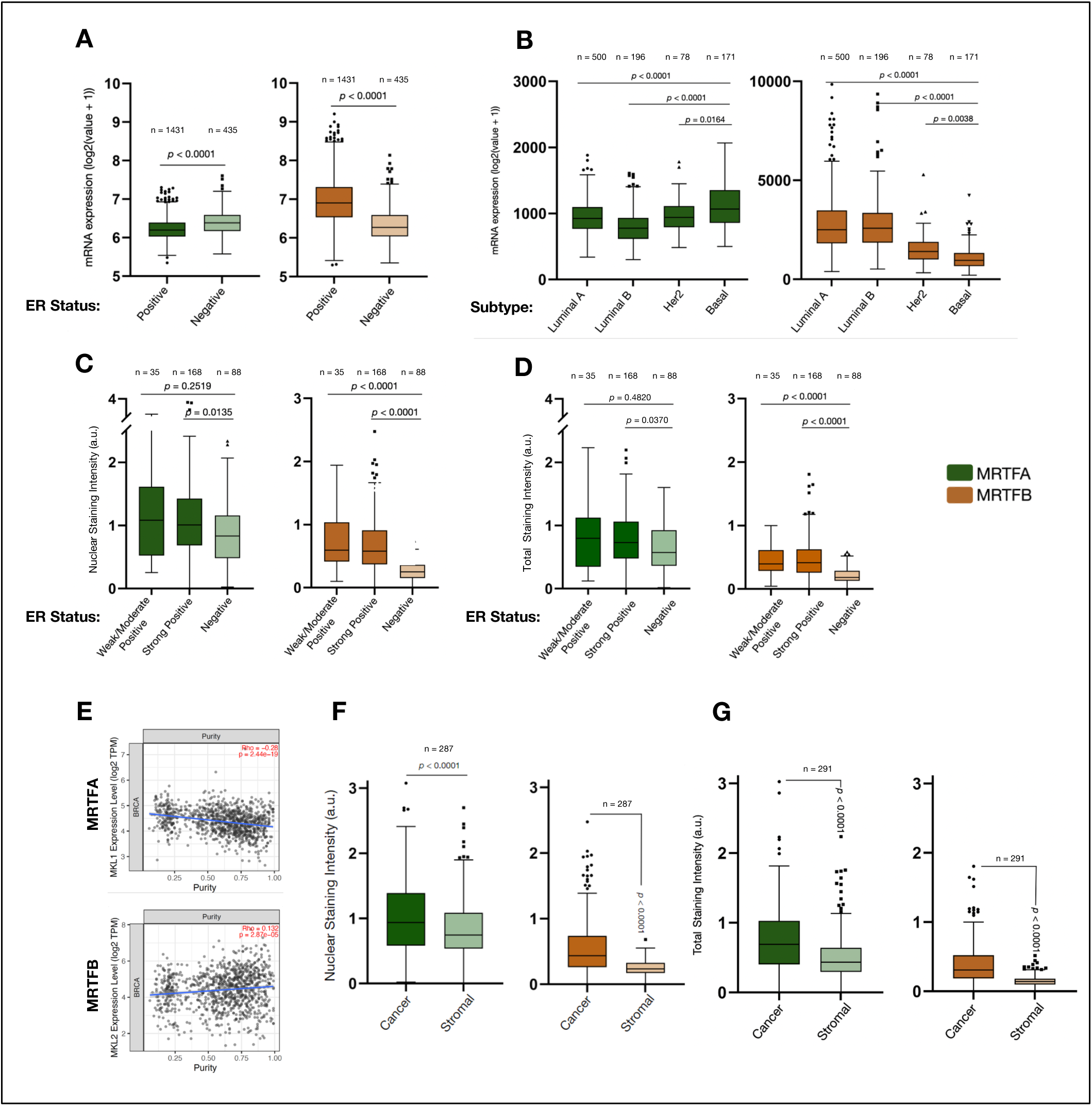
MRTFA and MRTFB have specific expression patterns based on ER status. (A) Box plots (Tukey presentation) for mRNA expression of MRTFA and MRTFB among estrogen receptor-positive (n = 1431) and estrogen receptor-negative (n = 435) invasive breast cancer cases. Data accessed via Molecular Taxonomy of Breast Cancer International Consortium (METABRIC) and Mann-Whitney test used to calculate *p*-values. (B) Box plots (Tukey presentation) for mRNA expression of MRTFA and MRTFB among Luminal A (n = 500), Luminal B (n = 196), Her2 (n = 78) and Basal (n = 171) molecular subtypes of breast cancer. Data accessed via The Cancer Genome Atlas (TCGA) and Kruskal-Wallis test used to calculate *p*-values. (C) Box plots (Tukey presentation) for nuclear staining intensity and total staining intensity (D) of MRTFA and MRTFB in panCK-positive tumor cells of primary tumor cores. Immunohistochemistry was used to identify tumors’ estrogen-receptor status as weak or moderate positive (1-10% and 11-50% staining, n = 35), strong positive (51-100% staining, n = 168), or negative (0% staining, n = 88). Kruskal-Wallis test used to calculate *p*-values. (E) Dot plot of all TCGA invasive breast cancer samples’ tumor purity as calculated by the TIMER algorithm as a function of *MRTFA* (*MKL1*) and *MRTFB* (*MKL2*) gene expression. (F) Box plots (Tukey presentation) for nuclear staining intensity and total staining intensity (G) of MRTFA and MRTFB in cancer and stromal compartments of primary tumor cores. Mann-Whitney test used to calculate *p*-values.

Based on the prominent expression of MRTFA in the TME, we decided to identify which stromal and immune cells expressed MRTFA/B in human breast cancers. Therefore, we queried an existing single cell sequencing database that contained 100,064 cells from 26 different breast cancer patients^32,33^. We found that MRTFB expression was highest in ACKR1+ endothelial cells and it was detected in approximately 27%-46% of various endothelial cell populations (**Fig. 5A**). Given the pathological roles of MRTFA in CAFs, we anticipated MRTFA to be most strongly expressed by the CAFs. Surprisingly, however, MRTFA expression was highest in the dendritic cells (DCs) in comparison to all the other cells of the TME. By using the TIMER2.0 algorithm on the TCGA dataset, we found that MRTFA, but not MRTFB, expression correlated strongly with DC infiltration and the level of correlation was higher for than the correlation between MRTFA expression and CAF infiltration (**Fig. 5B**). Next, we co-stained a breast cancer sample with MRTFA, panCK (tumor marker), CD31 (endothelial cell marker) and HLA-DRA, which is a major histocompatibility complex II (MHC II) protein that broadly marks most antigen presenting cells (APCs), including B-cells and DCs. We identified several HLA-DRA+ cells across the tumor microenvironment, but most prominently HLA-DRA+ cells showed a significant levels of MRTFA near expression CD31+ blood vessels (**Fig. 5C**). Since accumulation of perivascular APCs in human breast cancer has been recently associated with immune suppression and with DCIS progression to IBC, we decided to also stain the CHTN_BrCaProg3 TMA with our panel of antibodies^10^. Strikingly, we found strong MRTFA expression in the perivascular HLA-DRA+ cells in a DCIS sample (**Fig. 5D**). However, there was no prominent MRTFA expression in the HLA- DRA+ B-cell zones in the lymph nodes (**Fig. 5E**). Taken together, these data suggest that MRTFA may uniquely regulate antigen presenting APCs in the breast cancer TME.

**Figure 5.**
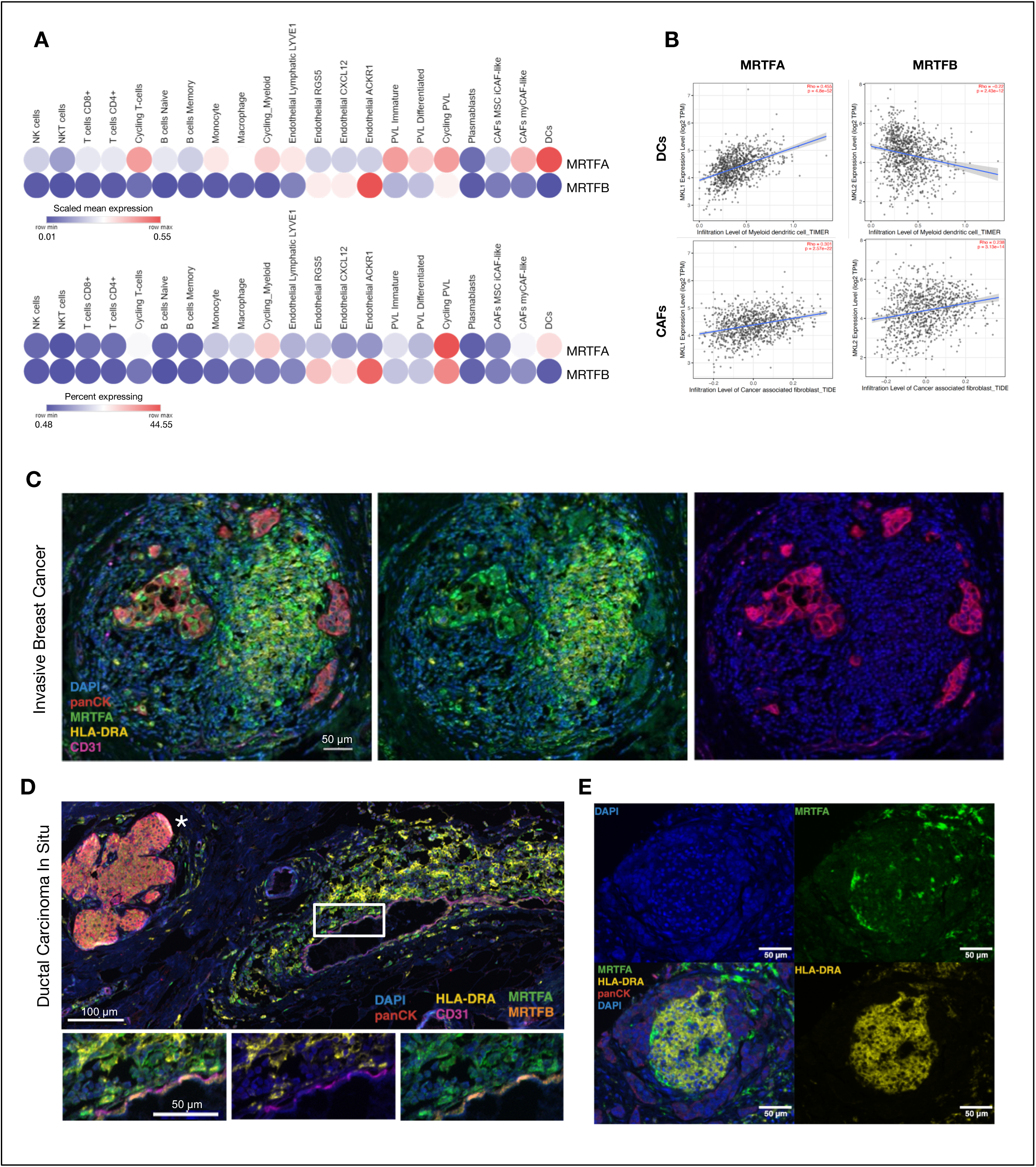
MRTFA shows a distinct expression pattern in antigen presenting cells in the tumor microenvironment. (A) Single-cell transcriptomic data showing scaled mean expression of and percent of cells expressing MRTFA and MRTFB in immune, structural, and antigen-presenting cell types of the tumor microenvironment (n = 26). Data accessed via Broad Institute Single Cell Portal. (B) Dendritic Cell (DC) and cancer associated fibroblast (CAF) enrichment scores based on TIMER algorithm as a function of *MRTFA* and *MRTFB* expression without tumor purity adjustments. (C-E) Multiplex image of (C) an invasive breast cancer sample from one patient included in the BRWG UIC-001-TMA (Scale bar, 50µm) and (D) in a ductal carcinoma *in situ* sample (Scale bar, 100µm for large view and scale bar, 50µm for insets) and (E) in a lymph node metastasis sample from CHTN_BrCaProg3 TMA showing cancer cells (panCK, red), endothelial cells (CD31, magenta), antigen presenting cells (HLA-DRA, yellow) and MRTFA (green) and MRTFB (cyan). (E) * shows the nearby DCIS lesion, rectangle shows the area of inset for the bottom panels.

The unique expression of MRTFA in APCs and its elevated expression in cancer cells of African American patients in our BRWG-UIC-001 TMA encouraged us to examine the potential connection between MRTFA expression and immune dysfunction, which is prominent in African American patients. First, we investigated how MRTFA could contribute to immune dysfunction. In non-small cell lung cancer and melanoma models MRTFA activity promotes programmed cell death 1 ligand (PD-L1) expression, which is an immune checkpoint protein that leads to cytotoxic T-cell exhaustion and tumor immune dysfunction^34–36^. Therefore, we interrogated the breast cancer single cell sequencing data for the expression of several well-established immune checkpoint proteins, such as PD-L1 (also known as CD274), PD-L2 (encoded by *PDCD1LG2*), V-set immunoregulatory receptor (VSIR, also known as V-domain immunoglobulin suppressor of T cell activation (VISTA) or C10orf54) and other immune suppressive molecules such as interleukin 10 (IL10) and Indoleamine 2,3-dioxygenase-1 (IDO1) (reviewed in 37). Among these immune suppressive molecules breast cancer DCs most prominently expressed VSIR (**Fig. 6A**). Interestingly, VSIR expression positively correlated with MRTFA expression in breast cancers in the TCGA dataset, to the same extent as some of the well-established direct transcriptional targets of MRTFA, such as myosin heavy chain and light chain genes, *MYH9* and *MYL9* (**Fig. 6B** and **Supplemental Fig. S4A** and **S4B**). To investigate whether MRTFA could directly regulate VSIR at the mRNA level, we used the University of California Santa Cruz (UCSC) Genome Browser to search experimentally validated SRF binding sites at the promoter region of the *VSIR* gene^38^. Indeed, we identified 5 distinct SRF binding sites that were experimentally validated by 12 different chromatin immunoprecipitation (ChiP) studies (**Fig. 6C**). In contrast, PDL1 contained only 2 SRF binding sites that were validated in 2 different experiments and PDL1 expression did not positively correlate with MRTFA expression in TCGA (**Supplemental Fig. S4C, S4D**). Taken together, these results suggest that MRTFA expression in APCs may lead to immune dysfunction by promoting VSIR expression.

**Figure 6.**
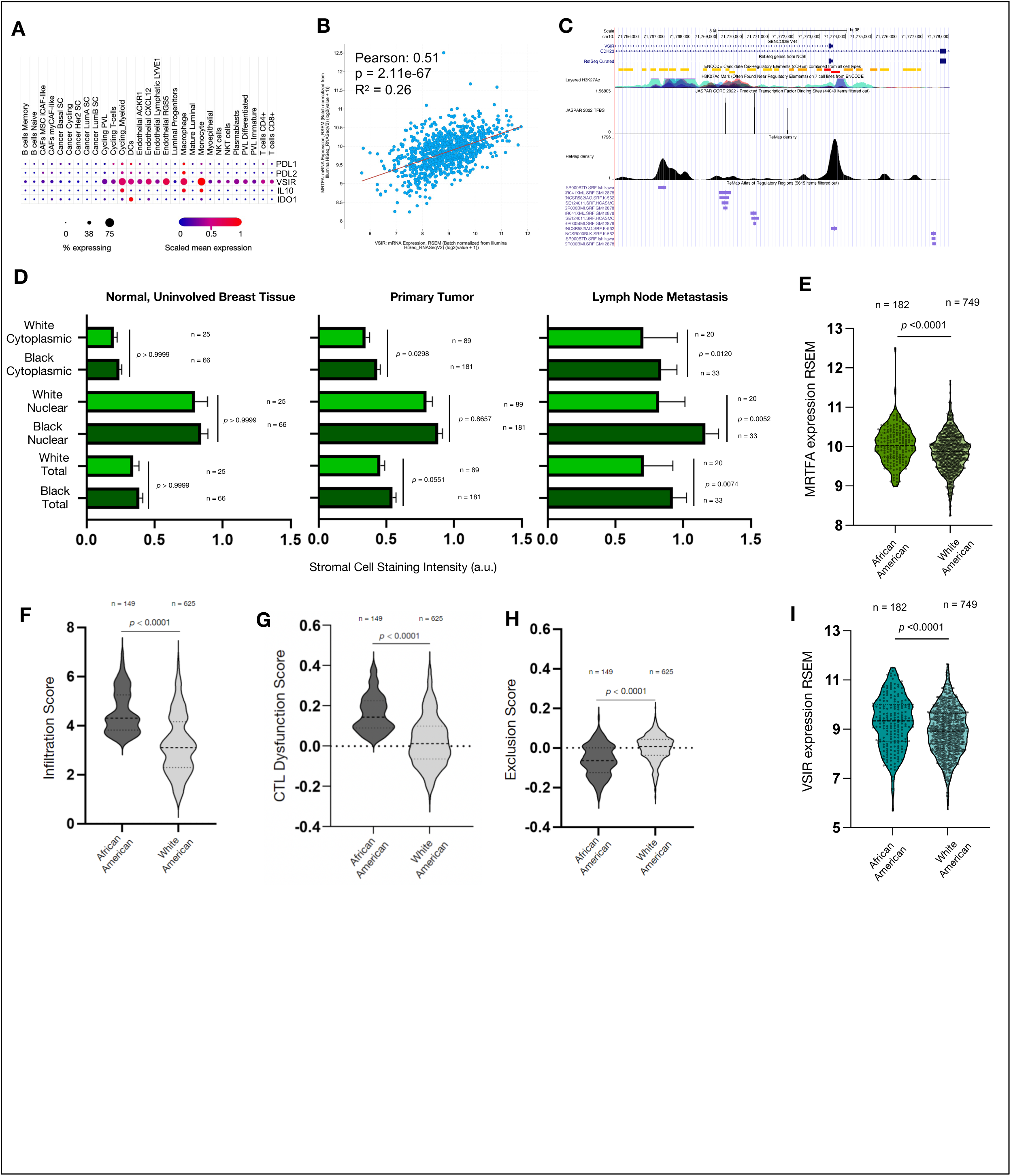
Immune exhaustion and MRTFA expression are elevated in tumors and the stroma of African American breast cancer patients. (**A**) Single cell RNA sequencing data showing expression of several immune suppressor protein. Data accessed via Broad Institute Single Cell Portal (**B**) Correlation between *MRTFA* and *VSIR* gene expression in TCGA invasive breast cancer dataset. (**C**) Transcription factor density plots showing SRF binding sites in the promoter region of human *VSIR* gene. Histone 3 Lysine 27 acetylation (H3K27Ac) show accessibility of chromatin regions, red, orange and yellow blocks in candidate cis-regulatory elements (cCREs) show promoter, and proximal cis enhancers respectively, JASPAR peaks show predicted SRF binding sites, ReMap density shows experimentally validated SRF binding sites by ChIP in experiments listed in purple font. Data accessed via http://genome.ucsc.edu. Session URL: https://genome.ucsc.edu/cgi-bin/hgTracks?db=hg38&lastVirtModeType=default&lastVirtModeExtraState=&virtModeType=default&virtMode=0&nonVirtPosition=&position=chr10%3A71761034%2D71778343&hgsid=1848347878_DsujX4MqToOVpoNLPdbawFAGvIeA. (**D**) MRTFA staining intensity of stromal (panCytokeratin negative) cells in the indicated selection of cores including normal/uninvolved breast tissue, primary tumor cores and lymph node metastasis cores in the BRWG UIC-001-TMA. *P-*values calculated by Mann-Whitney test. (**E**) MRTFA expression values (**F**) Cytotoxic T-cell Dysfunction (**G**) T-cell Exclusion and (**H**) T-cell infiltration scores and (**I**) VSIR expression values of breast cancer patients’ tumors in TCGA separated by race. *P-*values are calculated by Mann-Whitney test. (**B,E-I**) Data accessed via cBioportal.

Finally, we interrogated whether MRTFA could contribute to cancer health disparities by promoting immune dysfunction in African American patients. Social and racial injustices African American women endure are the root cause of immune weathering and dysfunction and they significantly contribute to postpartum lethality of mothers and to manifestation of chronic inflammatory diseases^7,39,40^. To measure the contribution of MRTFA expression to immune dysfunction, we measured MRTFA expression in African American patients in the stromal cells of the BRWG-UIC-001 TMA and in the TCGA dataset. We found that MRTFA was more highly expressed in stromal cells of lymph node metastases of the African American patients in the BRWG-UIC-001 TMA and in TCGA samples (**Fig. 6D and Fig. 6E**). Since elevated MRTFA expression in African American patients could be captured in the TCGA dataset, we tested whether African American breast cancer patients in TCGA also presented with a higher degree of immune dysfunction. Indeed, African American patients had elevated levels of T-cell inflitration, T-cell dysfunction as measured by the Tumor Immune Dysfunction and Exclusion (TIDE) algorithm^41^. Importantly, *VISR* gene expression was also modestly, but significantly increased in tumor samples from African American patients (**Fig. 6I**). Taken together, this data affirms the immune suppressive role of MRTFA expression in the TME and points to its activation as a biological mechanism contributing to breast cancer health disparities.

## Discussion

In this study, we used a series of multiplex imaging methods to analyze MRTFA/B expression in 2 different TMAs. CHTN_BrCaProg3 allowed us to interrogate MRTFA/B expression throughout transformation, tumorigenesis, and metastasis, while BRWG UIC-001-TMA allowed us to reveal racial disparities in MRTFA/B expression in invasive breast cancers. While our analyses complement the existing experimental knowledge on the role of MRTFA/B in breast cancer evolution and metastasis, we also report several novel findings on MRTFA/B activity across different breast cancer subtypes, racial categories and in cells of the TME. Importantly, we cross-validated these new findings with bioinformatics analyses of existing single cell and bulk tumor transcriptomics data.

Our first finding relates to MRTFA/B expression and activation. In published *in vitro* 2D models and 3D models, MRTFA/B primarily localizes to the cytoplasm unless it is stimulated by exogenous mitogens or cell adhesion to the extracellular matrix. This was in stark contrast to the human cancer samples we stained, where most tumor cells showed, on average, a higher degree of nuclear localization of MRTFA/B than their cytoplasmic localization. Importantly, however, MRTFA was not universally expressed in all DCIS or IBC samples: There were several samples that expressed very low levels of MRTFA/B, while a very few others had more abundant cytoplasmic localization than nuclear localization. Nevertheless, the observation that MRTFA/B show strong nuclear localization *in vivo*, as opposed to *in vitro*, suggests that the *in vitro* culture conditions may be lacking MRTFA/B stimulating factors that are present *in vivo*. One possibility is that the ECM composition, structure, and physical characteristics in tumor tissues may be more stimulatory toward MRTFA/B than the ECM *in vitro*. However, we reason that if ECM was the only source of MRTFA/B activation *in vivo*, we would have observed the most prominent MRTFA/B nuclear localization in the cells at the invasive front, which most extensively interact with ECM^27,42^. However, our imaging did not show as profound of a phenotype as is seen for other markers of the leading-edge such as p63 and Keratin 14 (Ref. 27). Therefore, we speculate that the broad nuclear MRTFA/B localization in human tissues may stem from cytokines and chemokines that are expressed by the stromal and immune cells of the TME. The identities and the prominent sources of these secreted factors will be the focus of future studies.

Our results also revealed differences in MRTFA/B expression across primary and metastatic tumors, different histological subtypes, and patients of different races. Molecular mechanisms that govern hematogenous and lymphatic metastases are distinct, and one limitation of our study is that we worked with only lymph node metastases^28,43^. Regardless of the dissemination route, expression of proteins that are important for metastasis are thought to be either (1) retained, (2) enriched or (3) lost at secondary sites. The third category includes, for example, regulators of epithelial to mesenchymal transition (EMT) that promote primary dissemination but become restrictive to outgrowth later in metastatic colonization^11^. Similarly, we had recently described that MRTFA/B are needed for the metastatic cascade, but their expression also presents with an immune vulnerability during colonization^16^. Therefore, we had speculated that MRTFA/B expression could be lost at the lymph node metastases even if the primary tumor expressed them. In contrast, we found that active MRTFA/B expression in the nucleus in the primary tumor cells positively correlated with lymph node expression. We also had anticipated that highest MRTFA expression levels would be observed in ER- negative subtype because single nucleotide polymorphisms within *MRTFA* gene have been associated with triple negative breast cancer risk^44,45^. However, race, but not ER status, was the strongest contributor to the MRTFA expression. The revelation that the tumors from African American patients expressed elevated levels of MRTFA led us to interrogate whether MRTFA+ metastatic tumor cells thrived due to an immune exhausted TME in these patients.

The TME of African American and White American breast patients present with quantitative differences in inflammatory signaling, macrophage infiltration and cytotoxic T-cell exhaustion^46–48^. Accordingly, we also found that the breast tumors from African American patients in TCGA presented with a higher degree of cytotoxic lymphocyte infiltration, but these tumors also had a higher level of exhaustion compared to breast tumors from White American patients. Our results are in line with comprehensive studies that show cytotoxic T-cell exhaustion in African American breast cancer patients^46,49^, and we speculate that this exhausted immune microenvironment contributes to immune evasion of aggressively metastatic MRTFA+ breast cancer cells in African American patients.

These observations suggest that there may be multiple therapeutic strategies to alleviate breast cancer disparities for African American breast cancer patients. One approach could be to target aggressively metastatic MRTFA+ cells through the use of immune checkpoint inhibitors as would be predicted from melanoma studies, where MRTFA’s transcriptional activity makes these cancer cells more vulnerable to anti-PD1 treatment through mechanosurveillance^16^. Another approach could be to target MRTFA+ APCs in the TME. Our studies revealed MRTFA+ APCs in the perivascular niche that resembled the perivascular immune suppressive APCs associated with DCIS progression to IBC^10^. Based on the positive correlation between MRTFA expression and the immune checkpoint protein VSIR expression, we propose that the use of anti-VSIR neutralizing antibodies can prevent the immune suppressive function of these MRTFA+ APCs and help alleviate cancer health disparities that African American breast cancer suffer from^50^. The overall increase in MRTFA expression in the TME of African American patients’ breast tumors also point to a broader role of immune suppression by MRTFA expressing stromal cells such as those residing in tertiary lymphoid structures (TLS) or the recently described “suppressed expansion” zones, or the CAFs ^51,52^. We also consider that some of the immune suppressive functions of MRTFA could be attributed to its negative role in dendritic cell maturation as has been described in recent murine models of MRTFA knockout or B2 Integrin-Kindlin3 loss-of-function^53,54^. Future work will investigate how pharmacological suppression of MRTFA in patients would alter the tumor immune microenvironment to help alleviate breast cancer disparities.

## Acknowledgements

The authors wish to acknowledge the Breast Cancer Working Group and Translational Pathology Shared Resource of the University of Illinois Cancer Center, and the University of Illinois Biorepository, for providing access to the tissue microarrays. We thank Eileen Brister and Ryan Deaton from the Research Histology and Imaging Collaborative Core for their assistance with automated tissue staining, image acquisition and implementation of HaLO segmentation and measurements; Klara Valyi-Nagy for their UI Health Biorepository access, Peter Gann, for their help with multiplex antibody panel design and thoughtful discussions. Research reported in this publication was supported in part by the University of Illinois Cancer Center Biostatistics Shared Resource (BSR) core.

## Author contributions

S.W. and E.E.E. initiated the project, wrote the manuscript, analyzed data; S.W. collected data; K.L., A.G., M.H., conducted experiments, collected data, edited the manuscript; K.F.H. collected data, provided study materials, critically reviewed the manuscript drafts; E.L.W. selected tissue samples to be tested, curated and selected tissue samples for BRWG UIC-001-TMA construction; V.M. constructed the BRWG UIC-001-TMA; Z.C. assisted with statistical analyses and data interpretation; X.L. and X.W conducted the bioinformatics analyses for T-cell dysfunction; M.S. optimized multiplex staining, collected and interpreted data.

---

**Supplemental Figure 1. Related to Figure 2.**
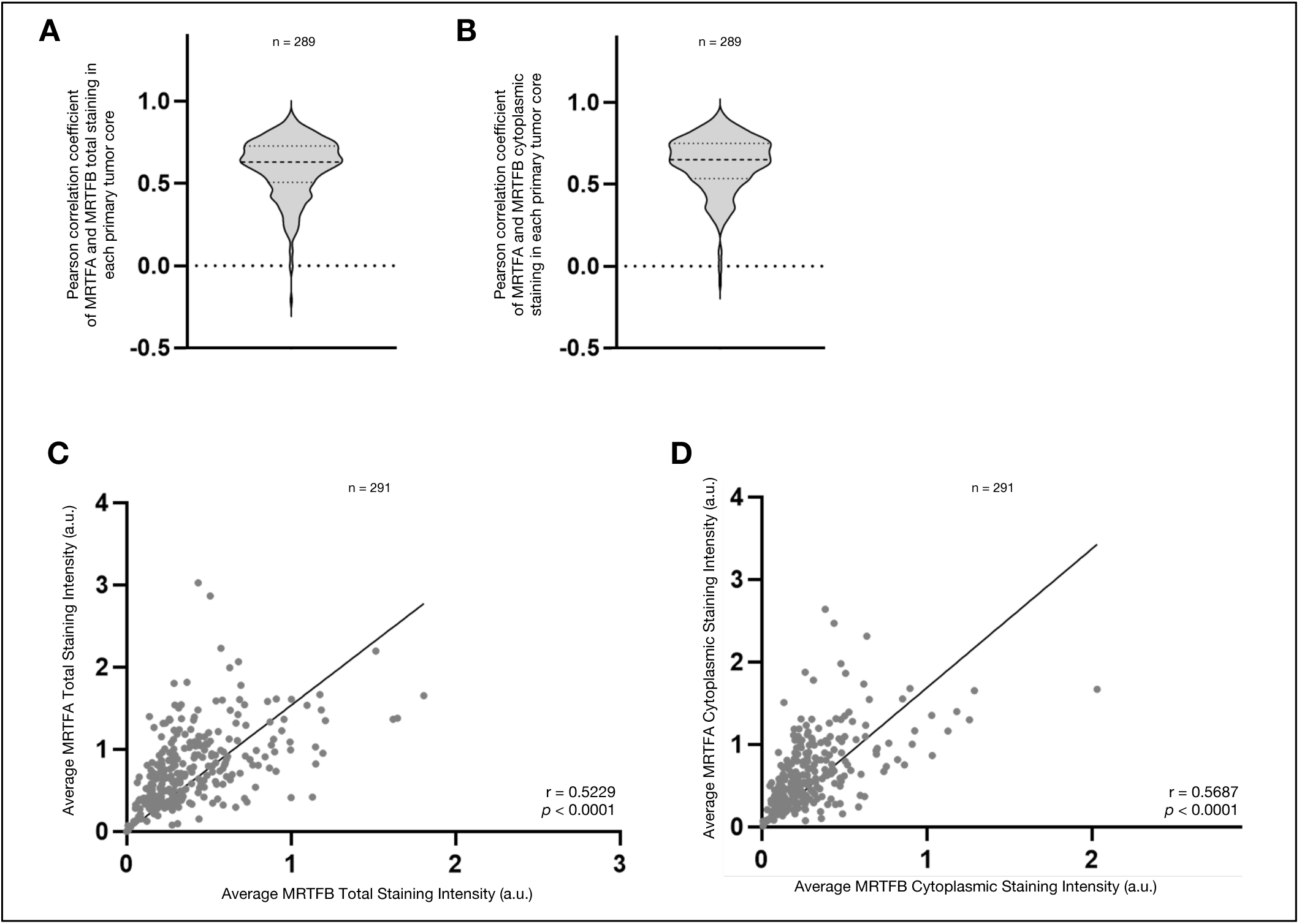
(A) Violin plot showing distribution of Pearson correlation coefficients of average total MRTFA and MRTFB staining intensities and average cytoplasmic MRTFA and MRTFB staining intensities (B) in panCK-positive tumor cells of primary tumor cores (n = 289). Mann-Whitney test used to calculate *p*-values. (C) Pearson correlation plot and simple linear regression of average total MRTFA and MRTFB staining intensities and average cytoplasmic MRTFA and MRTFB staining intensities (D) in panCK-positive tumor cells of primary tumor cores (n = 291).

**Supplemental Figure 2. Related to Figure 3.**
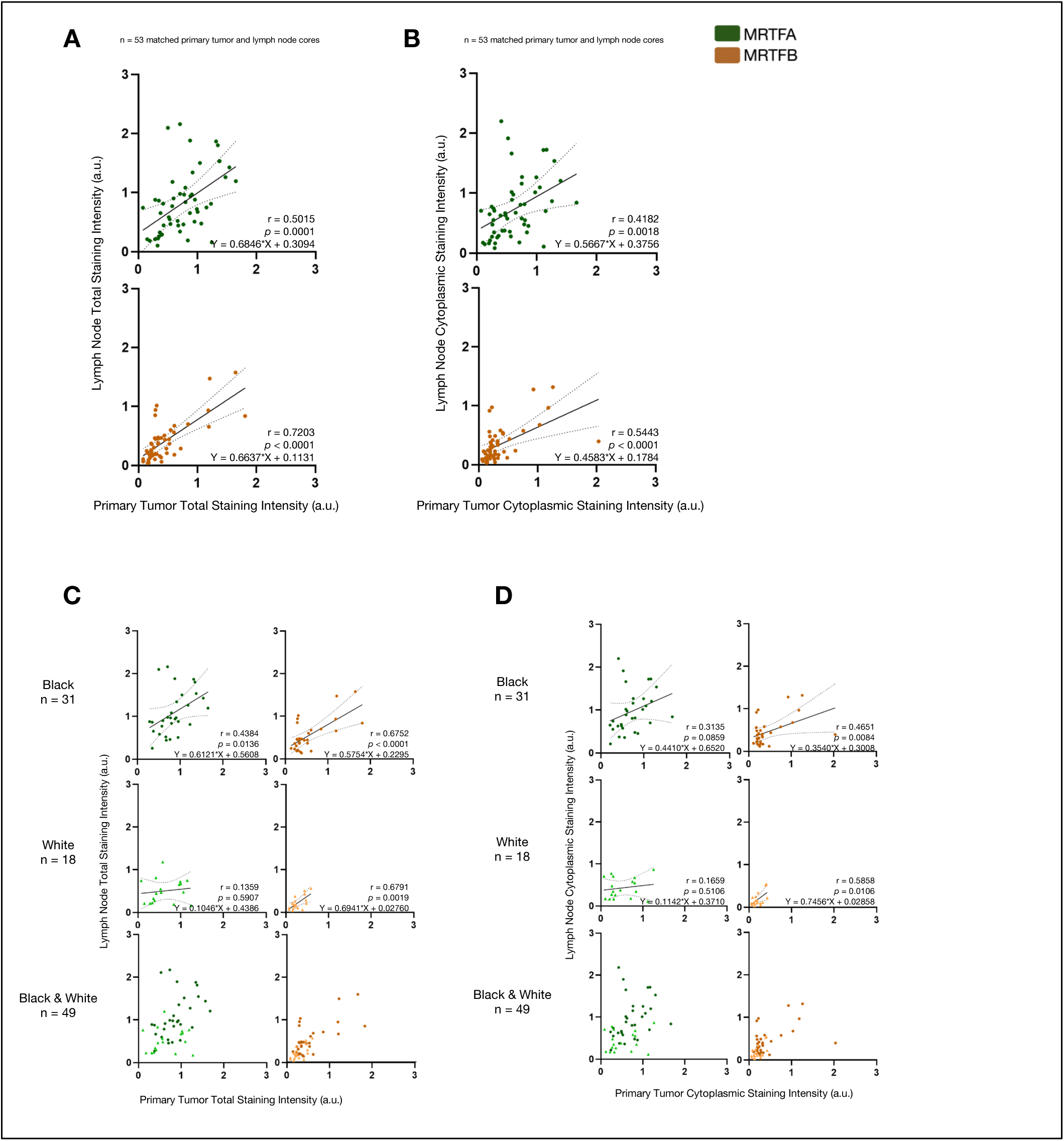
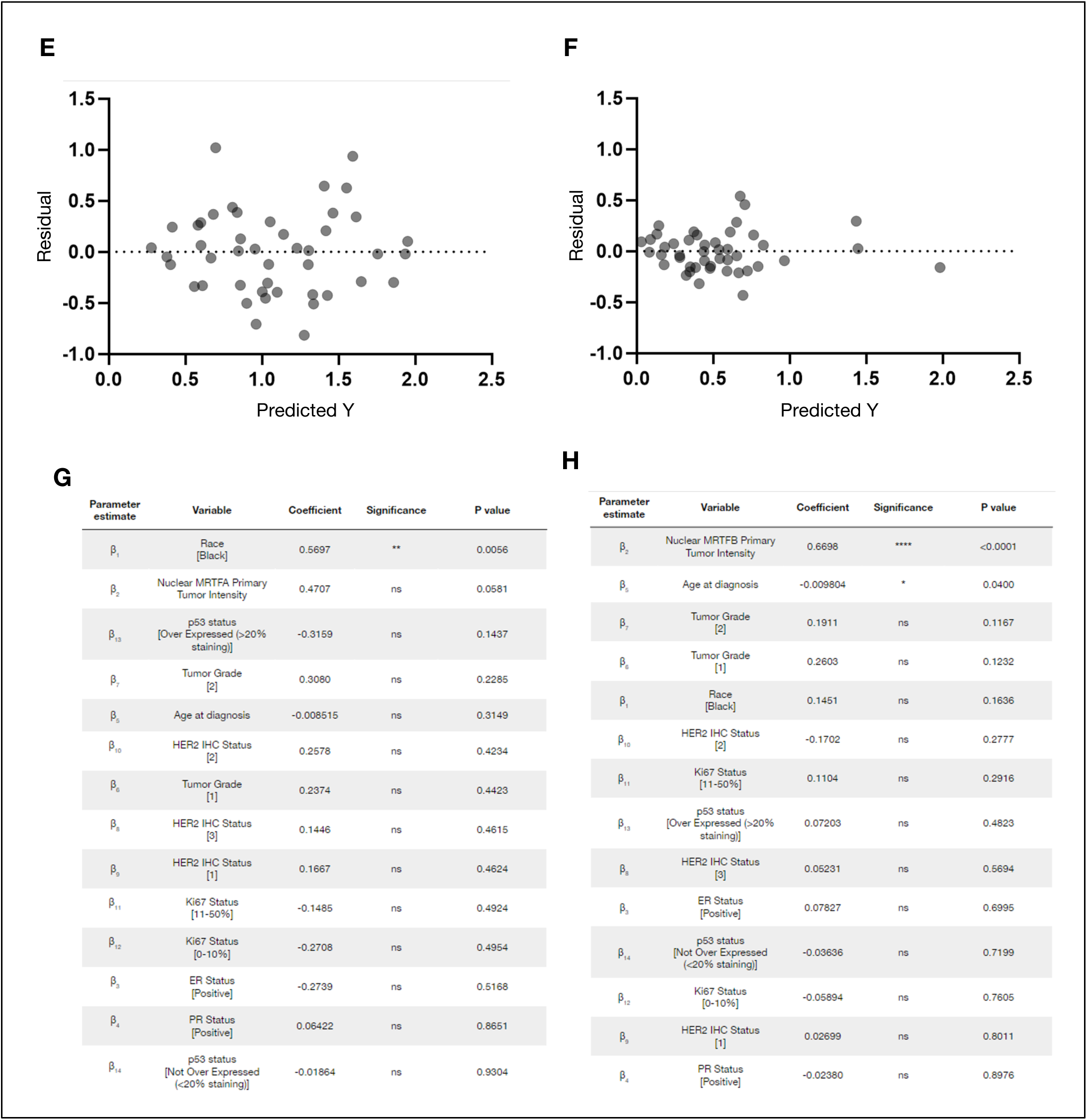
(A) Pearson correlation plots of primary tumor total staining intensity and matching lymph node metastasis total staining intensity of MRTFA and MRTFB. Dotted lines signify 99% confidence interval (n = 53). (B) Pearson correlation plots of primary tumor cytoplasmic staining intensity and matching lymph node metastasis cytoplasmic staining intensity of MRTFA and MRTFB. Dotted lines signify 99% confidence interval (n = 53). (C) Pearson correlation plots of primary tumor total staining intensity and matching lymph node total staining intensity of MRTFA and MRTFB for self-identified Black/African American patients (n = 31) and White patients (n = 18). Dotted lines signify 99% confidence interval. (D) Pearson correlation plots of primary tumor cytoplasmic staining intensity and matching lymph node cytoplasmic staining intensity of MRTFA and MRTFB for self-identified Black/African American patients (n = 31) and White patients (n = 18). Dotted lines signify 99% confidence interval. (E) Residual plot for multiple linear regression analysis of nuclear MRTFA lymph node intensity with patient demographic and tumor characteristics predictor variables. (F) Residual plot for multiple linear regression analysis of nuclear MRTFB lymph node intensity with patient demographic and tumor characteristics predictor variables. (G) Complete multiple linear regression results with all predictor variables for nuclear MRTFA lymph node intensity organized by P-value. (H) Complete multiple linear regression results with all predictor variables for nuclear MRTFB lymph node intensity organized by P-value. Significance level for *p*-values: * (*p*<0.05), ** (*p*<0.01), *** (*p*<0.001), **** (*p*<0.0001).

**Supplemental Figure 3. Related to Figure 4.**
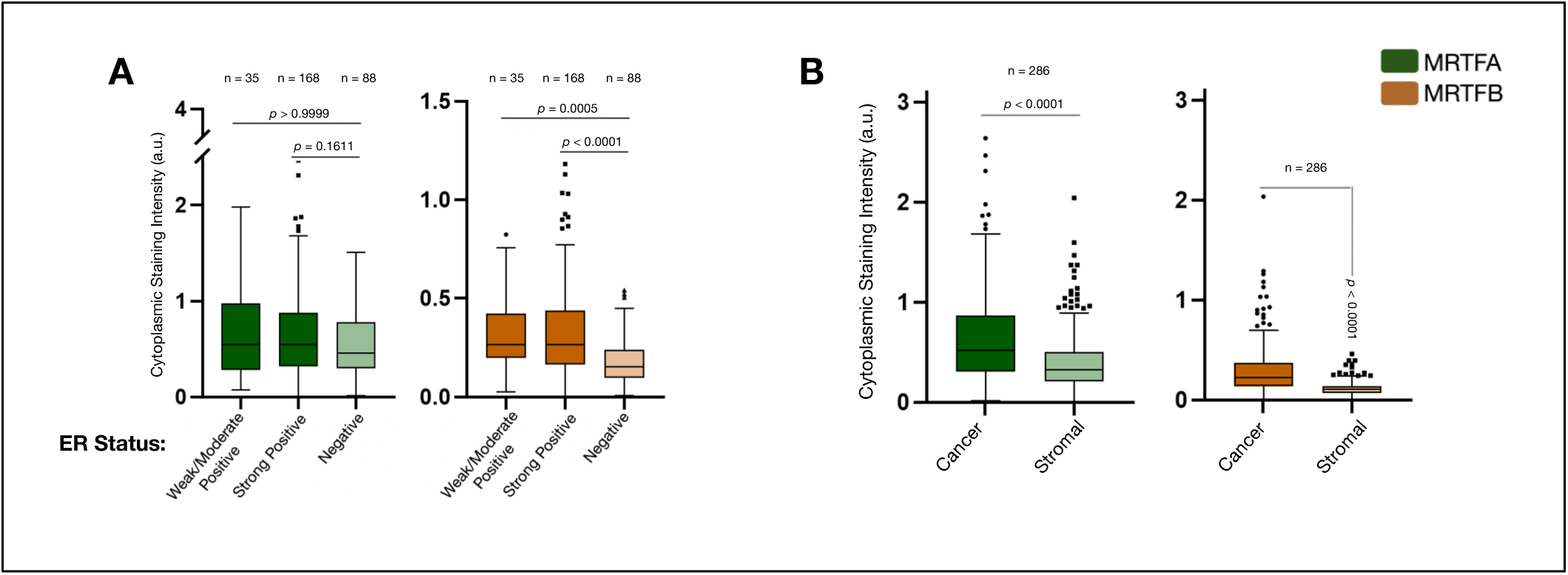
(A) Box plots (Tukey presentation) for cytoplasmic staining intensity of MRTFA and MRTFB in panCK-positive tumor cells of primary tumor cores. Immunohistochemistry was used to identify tumors’ estrogen-receptor status as weak or moderate positive (1-10% and 11-50% staining, n = 35), strong positive (51-100% staining, n = 168), or negative (0% staining, n = 88). Kruskal-Wallis test used to calculate *p*-values. (B) Box plots (Tukey presentation) for cytoplasmic staining intensity of MRTFA and MRTFB in cancer and stromal compartments of primary tumor cores. Mann-Whitney test used to calculate *p*-values.

**Supplemental Figure 4. Related to Figure 6.**
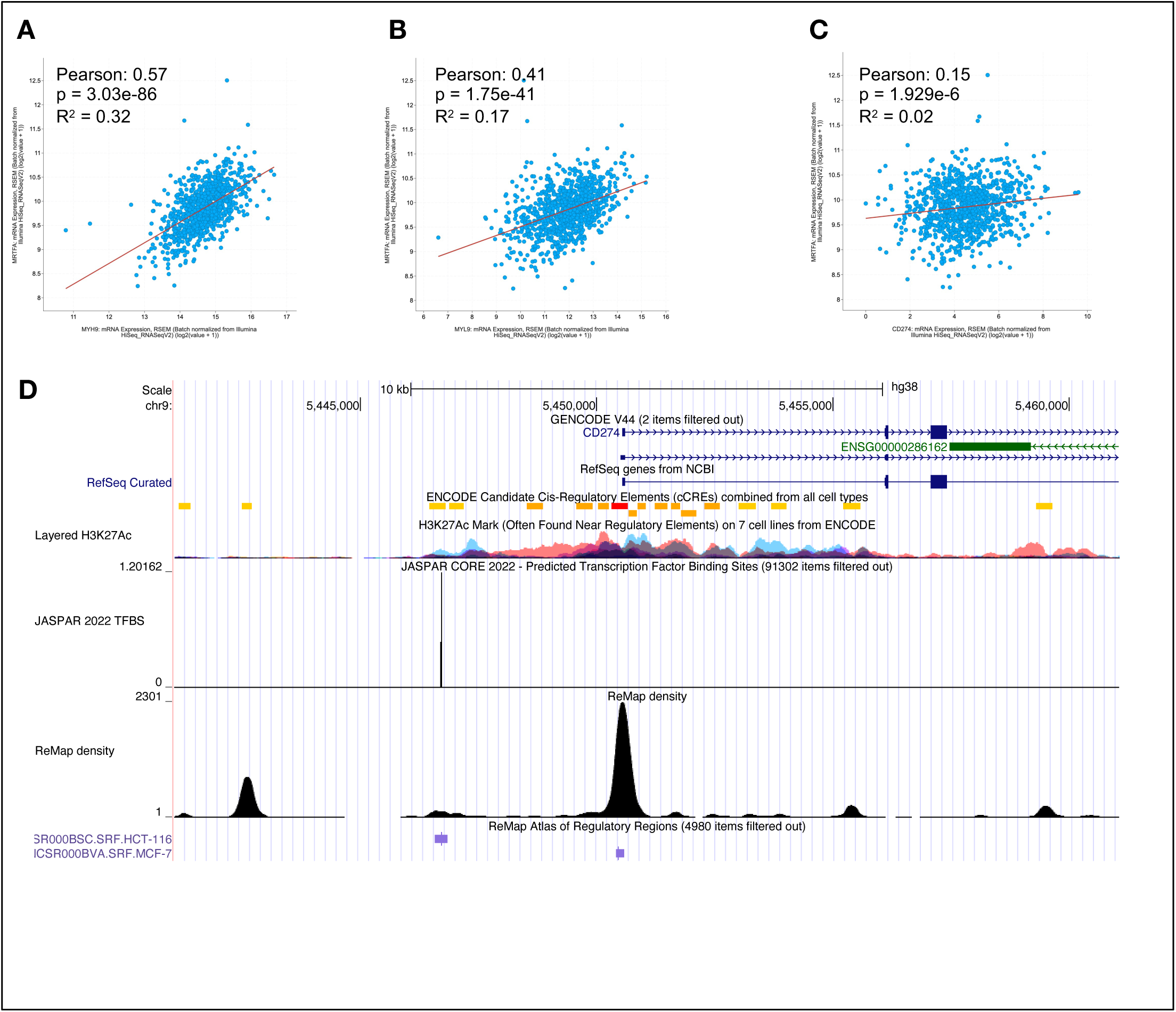
(**A-C**) Correlation between *MRTFA* expression and (**A**) *MYH9* (**B**) *MYL9* and (**C**) PDL1 (encoded by *CD274*) gene expression in TCGA invasive breast cancer dataset. Data accessed via cBioportal. (**D**) Transcription factor density plots showing SRF binding sites in the promoter region of human *CD274* gene. Histone 3 Lysine 27 acetylation (H3K27Ac) show accessibility of chromatin regions, red, orange and yellow blocks in candidate cis-regulatory elements (cCREs) show promoter, and proximal cis enhancers respectively, JASPAR peaks show predicted SRF binding sites, ReMap density shows experimentally validated SRF binding sites by ChIP in experiments listed in purple font. Data accessed via http://genome.ucsc.edu. Session URL: https://genome.ucsc.edu/cgi-bin/hgTracks?db=hg38&lastVirtModeType=default&lastVirtModeExtraState=&virtModeType=default&virtMode=0&nonVirtPosition=&position=chr9%3A5445502%2D5455508&hgsid=1848344652_Gzjysw7sxcCdpuiGrb4ZWbQgKRml.

## References

1 Giaquinto, A. N. et al. Breast Cancer Statistics, 2022. CA Cancer J Clin 72, 524–541, doi:10.3322/caac.21754 (2022).

2 Hoskins, K. F., Danciu, O. C., Ko, N. Y. & Calip, G. S. Association of Race/Ethnicity and the 21-Gene Recurrence Score With Breast Cancer-Specific Mortality Among US Women. JAMA Oncol 7, 370–378, doi:10.1001/jamaoncol.2020.7320 (2021).

3 Alvarez, A., Bernal, A. M. & Anampa, J. Racial disparities in overall survival after the introduction of cyclin-dependent kinase 4/6 inhibitors for patients with hormone receptor-positive, HER2-negative metastatic breast cancer. Breast Cancer Res Treat 198, 75–88, doi:10.1007/s10549-022-06847-2 (2023).

4 Linnenbringer, E., Gehlert, S. & Geronimus, A. T. Black-White Disparities in Breast Cancer Subtype: The Intersection of Socially Patterned Stress and Genetic Expression. AIMS Public Health 4, 526–556, doi:10.3934/publichealth.2017.5.526 (2017).

5 Carlos, R. C. et al. Linking Structural Racism and Discrimination and Breast Cancer Outcomes: A Social Genomics Approach. J Clin Oncol 40, 1407–1413, doi:10.1200/JCO.21.02004 (2022).

6 Islami, F. et al. American Cancer Society’s report on the status of cancer disparities in the United States, 2021. CA Cancer J Clin 72, 112–143, doi:10.3322/caac.21703 (2022).

7 Valencia, C. I., Gachupin, F. C., Molina, Y. & Batai, K. Interrogating Patterns of Cancer Disparities by Expanding the Social Determinants of Health Framework to Include Biological Pathways of Social Experiences. Int J Environ Res Public Health 19, doi:10.3390/ijerph19042455 (2022).

8 Wiechmann, L. & Kuerer, H. M. The molecular journey from ductal carcinoma in situ to invasive breast cancer. Cancer 112, 2130–2142, doi:10.1002/cncr.23430 (2008).

9 Gibson, S. V. et al. Everybody needs good neighbours: the progressive DCIS microenvironment. Trends Cancer 9, 326–338, doi:10.1016/j.trecan.2023.01.002 (2023).

10 Risom, T. et al. Transition to invasive breast cancer is associated with progressive changes in the structure and composition of tumor stroma. Cell 185, 299–310 e218, doi:10.1016/j.cell.2021.12.023 (2022).

11 Er, E. E., Tello-Lafoz, M. & Huse, M. Mechanoregulation of Metastasis beyond the Matrix. Cancer Res 82, 3409–3419, doi:10.1158/0008-5472.CAN-22-0419 (2022).

12 Er, E. E. et al. Pericyte-like spreading by disseminated cancer cells activates YAP and MRTF for metastatic colonization. Nat Cell Biol 20, 966–978, doi:10.1038/s41556-018-0138-8 (2018).

13 Kim, T. et al. MRTF potentiates TEAD-YAP transcriptional activity causing metastasis. EMBO J 36, 520–535, doi:10.15252/embj.201695137 (2017).

14 Medjkane, S., Perez-Sanchez, C., Gaggioli, C., Sahai, E. & Treisman, R. Myocardin-related transcription factors and SRF are required for cytoskeletal dynamics and experimental metastasis. Nat Cell Biol 11, 257–268, doi:10.1038/ncb1833 (2009).

15 Seifert, A. & Posern, G. Tightly controlled MRTF-A activity regulates epithelial differentiation during formation of mammary acini. Breast Cancer Res 19, 68, doi:10.1186/s13058-017-0860-3 (2017).

16 Tello-Lafoz, M. et al. Cytotoxic lymphocytes target characteristic biophysical vulnerabilities in cancer. Immunity 54, 1037–1054 e1037, doi:10.1016/j.immuni.2021.02.020 (2021).

17 Li, S., Chang, S., Qi, X., Richardson, J. A. & Olson, E. N. Requirement of a myocardin-related transcription factor for development of mammary myoepithelial cells. Mol Cell Biol 26, 5797–5808, doi:10.1128/MCB.00211-06 (2006).

18 Li, J. et al. Myocardin-related transcription factor B is required in cardiac neural crest for smooth muscle differentiation and cardiovascular development. Proc Natl Acad Sci U S A 102, 8916–8921, doi:10.1073/pnas.0503741102 (2005).

19 Wei, K., Che, N. & Chen, F. Myocardin-related transcription factor B is required for normal mouse vascular development and smooth muscle gene expression. Dev Dyn 236, 416–425, doi:10.1002/dvdy.21041 (2007).

20 Oh, J., Richardson, J. A. & Olson, E. N. Requirement of myocardin-related transcription factor-B for remodeling of branchial arch arteries and smooth muscle differentiation. Proc Natl Acad Sci U S A 102, 15122–15127, doi:10.1073/pnas.0507346102 (2005).

21 Weinl, C. et al. Endothelial SRF/MRTF ablation causes vascular disease phenotypes in murine retinae. J Clin Invest 123, 2193–2206, doi:10.1172/JCI64201 (2013).

22 Foster, C. T., Gualdrini, F. & Treisman, R. Mutual dependence of the MRTF-SRF and YAP-TEAD pathways in cancer-associated fibroblasts is indirect and mediated by cytoskeletal dynamics. Genes Dev 31, 2361–2375, doi:10.1101/gad.304501.117 (2017).

23 Lopez-Garcia, M. A., Geyer, F. C., Lacroix-Triki, M., Marchio, C. & Reis-Filho, J. S. Breast cancer precursors revisited: molecular features and progression pathways. Histopathology 57, 171–192, doi:10.1111/j.1365-2559.2010.03568.x (2010).

24 Miralles, F., Posern, G., Zaromytidou, A. I. & Treisman, R. Actin dynamics control SRF activity by regulation of its coactivator MAL. Cell 113, 329–342, doi:10.1016/s0092-8674(03)00278-2 (2003).

25 Vartiainen, M. K., Guettler, S., Larijani, B. & Treisman, R. Nuclear actin regulates dynamic subcellular localization and activity of the SRF cofactor MAL. Science 316, 1749–1752, doi:10.1126/science.1141084 (2007).

26 Montel, L., Sotiropoulos, A. & Henon, S. The nature and intensity of mechanical stimulation drive different dynamics of MRTF-A nuclear redistribution after actin remodeling in myoblasts. PLoS One 14, e0214385, doi:10.1371/journal.pone.0214385 (2019).

27 Cheung, K. J., Gabrielson, E., Werb, Z. & Ewald, A. J. Collective invasion in breast cancer requires a conserved basal epithelial program. Cell 155, 1639–1651, doi:10.1016/j.cell.2013.11.029 (2013).

28 Massague, J. & Obenauf, A. C. Metastatic colonization by circulating tumour cells. Nature 529, 298–306, doi:10.1038/nature17038 (2016).

29 Jehanno, C. et al. Nuclear translocation of MRTFA in MCF7 breast cancer cells shifts ERalpha nuclear/genomic to extra-nuclear/non genomic actions. Mol Cell Endocrinol 530, 111282, doi:10.1016/j.mce.2021.111282 (2021).

30 Kerdivel, G. et al. Activation of the MKL1/actin signaling pathway induces hormonal escape in estrogen-responsive breast cancer cell lines. Mol Cell Endocrinol 390, 34–44, doi:10.1016/j.mce.2014.03.009 (2014).

31 Livasy, C. A. et al. Phenotypic evaluation of the basal-like subtype of invasive breast carcinoma. Mod Pathol 19, 264–271, doi:10.1038/modpathol.3800528 (2006).

32 Wu, S. Z. et al. A single-cell and spatially resolved atlas of human breast cancers. Nat Genet 53, 1334–1347, doi:10.1038/s41588-021-00911-1 (2021).

33 Tarhan, L., et al. Single Cell Portal: an interactive home for single-cell genomics data. bioRxiv, doi:10.1101/2023.07.13.548886 (2023).

34 Wei, S. C., Duffy, C. R. & Allison, J. P. Fundamental Mechanisms of Immune Checkpoint Blockade Therapy. Cancer Discov 8, 1069–1086, doi:10.1158/2159-8290.CD-18-0367 (2018).

35 Du, F. et al. MRTF-A-NF-kappaB/p65 axis-mediated PDL1 transcription and expression contributes to immune evasion of non-small-cell lung cancer via TGF-beta. Exp Mol Med 53, 1366–1378, doi:10.1038/s12276-021-00670-3 (2021).

36 Foda, B. M., Misek, S. A., Gallo, K. A. & Neubig, R. R. Inhibition of the Rho/MRTF pathway improves the response of BRAF-resistant melanoma to PD1/PDL1 blockade. bioRxiv, 2023.2012.2020.572555, doi:10.1101/2023.12.20.572555 (2023).

37 Wculek, S. K. et al. Dendritic cells in cancer immunology and immunotherapy. Nat Rev Immunol 20, 7–24, doi:10.1038/s41577-019-0210-z (2020).

38 Kent, W. J. et al. The human genome browser at UCSC. Genome Res 12, 996–1006, doi:10.1101/gr.229102 (2002).

39 Geronimus, A. T. The weathering hypothesis and the health of African-American women and infants: evidence and speculations. Ethn Dis 2, 207–221 (1992).

40 Das, A. How does race get “under the skin”?: inflammation, weathering, and metabolic problems in late life. Soc Sci Med 77, 75–83, doi:10.1016/j.socscimed.2012.11.007 (2013).

41 Jiang, P. et al. Signatures of T cell dysfunction and exclusion predict cancer immunotherapy response. Nat Med 24, 1550–1558, doi:10.1038/s41591-018-0136-1 (2018).

42 Han, Y. L. et al. Cell swelling, softening and invasion in a three-dimensional breast cancer model. Nat Phys 16, 101–108, doi:10.1038/s41567-019-0680-8 (2020).

43 Jones, D., Pereira, E. R. & Padera, T. P. Growth and Immune Evasion of Lymph Node Metastasis. Front Oncol 8, 36, doi:10.3389/fonc.2018.00036 (2018).

44 Purrington, K. S. et al. Genome-wide association study identifies 25 known breast cancer susceptibility loci as risk factors for triple-negative breast cancer. Carcinogenesis 35, 1012–1019, doi:10.1093/carcin/bgt404 (2014).

45 Lindstrom, S. et al. Genome-wide association study identifies multiple loci associated with both mammographic density and breast cancer risk. Nat Commun 5, 5303, doi:10.1038/ncomms6303 (2014).

46 Yao, S. et al. Breast Tumor Microenvironment in Black Women: A Distinct Signature of CD8+ T-Cell Exhaustion. J Natl Cancer Inst 113, 1036–1043, doi:10.1093/jnci/djaa215 (2021).

47 Martin, D. N. et al. Differences in the tumor microenvironment between African-American and European-American breast cancer patients. PLoS One 4, e4531, doi:10.1371/journal.pone.0004531 (2009).

48 Marczyk, M. et al. Tumor immune microenvironment of self-identified African American and non-African American triple negative breast cancer. NPJ Breast Cancer 8, 88, doi:10.1038/s41523-022-00449-3 (2022).

49 O’Meara, T. et al. Immune microenvironment of triple-negative breast cancer in African-American and Caucasian women. Breast Cancer Res Treat 175, 247–259, doi:10.1007/s10549-019-05156-5 (2019).

50 Martin, A. S. et al. VISTA expression and patient selection for immune-based anticancer therapy. Front Immunol 14, 1086102, doi:10.3389/fimmu.2023.1086102 (2023).

51 Schumacher, T. N. & Thommen, D. S. Tertiary lymphoid structures in cancer. Science 375, eabf9419, doi:10.1126/science.abf9419 (2022).

52 Danenberg, E. et al. Breast tumor microenvironment structures are associated with genomic features and clinical outcome. Nat Genet 54, 660–669, doi:10.1038/s41588-022-01041-y (2022).

53 Guenther, C. et al. A beta2-Integrin/MRTF-A/SRF Pathway Regulates Dendritic Cell Gene Expression, Adhesion, and Traction Force Generation. Front Immunol 10, 1138, doi:10.3389/fimmu.2019.01138 (2019).

54 Morrison, V. L. et al. Loss of beta2-integrin-mediated cytoskeletal linkage reprogrammes dendritic cells to a mature migratory phenotype. Nat Commun 5, 5359, doi:10.1038/ncomms6359 (2014).

